# Intersubject consistent dynamic connectivity during natural vision revealed by functional MRI

**DOI:** 10.1101/796433

**Authors:** Xin Di, Bharat B Biswal

## Abstract

The functional communications between brain regions are thought to be dynamic. However, it is usually difficult to elucidate whether the observed dynamic connectivity is functionally meaningful or simply due to noise during unconstrained task conditions such as resting-state. During naturalistic conditions, such as watching a movie, it has been shown that local brain activities, e.g. in the visual cortex, are consistent across subjects. Following similar logic, we propose to study intersubject correlations of the time courses of dynamic connectivity during naturalistic conditions to extract functionally meaningful dynamic connectivity patterns. We analyzed a functional MRI (fMRI) dataset when the subjects watched a short animated movie. We calculated dynamic connectivity by using sliding window technique, and quantified the intersubject correlations of the time courses of dynamic connectivity. Although the time courses of dynamic connectivity are thought to be noisier than the original signals, we found similar level of intersubject correlations of dynamic connectivity to those of regional activity. Most importantly, highly consistent dynamic connectivity could occur between regions that did not show high intersubject correlations of regional activity, and between regions with little stable functional connectivity. The analysis highlighted higher order brain regions such as the default mode network that dynamically interacted with posterior visual regions during the movie watching, which may be associated with the understanding of the movie.

**Highlights:**

- Intersubject consistency may provide a complementary approach to study brain dynamic connectivity
- Widespread brain regions showed highly consistent dynamic connectivity during movie watching, while these regions themselves did not show highly consistent regional activity
- Consistent dynamic connectivity often occurred between regions from different functional systems

## 1. Introduction

The functional communications between spatially remote brain regions, especially the dynamics of connectivity, is a key to understand brain functions (Bullmore and Sporns, 2012; Friston, 2011; Park and Friston, 2013). Recently, the dynamics of connectivity has drawn increasing interests of research, especially in resting-state (Allen et al., 2014; Fu et al., 2019, 2018; Hutchison et al., 2013). However, due to the unconstrained nature of resting-state, it is difficult to elucidate whether the observed changes of connectivity across sliding windows are due to real fluctuations of functional communications, or simply due to random fluctuations (Lindquist et al., 2014). Moreover, the blood-oxygen-level dependent (BOLD) signals measured by fMRI are sensitive to physiological noises, such as respiration, heartbeat (Teichert et al., 2010), and head motion (Power et al., 2012), which may give rise to spurious correlation estimates for short window.

One way to capture meaningful dynamic functional connectivity is to manipulate subjects’ mental states during the course of scan, so that there is known reference for the changes of connectivity. For example, in a typical task-based fMRI study with blocked design, different task conditions are assigned as blocks. Therefore, the time courses of dynamic connectivity can be correlated with the task design to identify task related connectivity changes (Di et al., 2015; Rosenthal et al., 2017). An alternative approach is to expose the subjects with naturalistic stimuli, such as a short movie. Although there is no predefined references of dynamic connectivity changes, one may take advantage of the phenomenon of intersubject correlation to capture changes that are consistent across different subjects (Hasson et al., 2004; Nastase et al., 2019). In the seminal study, Hasson and colleagues calculated intersubject correlations of the time series of BOLD signal (Figure 1A) when the subjects were watching a movie (Hasson et al., 2004). They demonstrated that several brain regions, especially the visual cortex, are highly correlated across subjects during the movie watching. We propose that similar approach can be applied to the time courses of dynamic connectivity to capture meaningful functional communication dynamics during natural vision. Specifically, dynamic connectivity is usually calculated using a sliding window approach, so that a time series of dynamic connectivity can be obtained. The time courses of dynamic connectivity can then be correlated across-subjects (Figure 1B). If the dynamic connectivity reflects real time functional communications between regions that are caused by the viewing of natural stimuli, then the time courses of dynamic connectivity from different subjects should somehow correlated. Therefore, we can apply intersubject correlation method to identify meaningful dynamic communications between regions.

**Figure 1.**
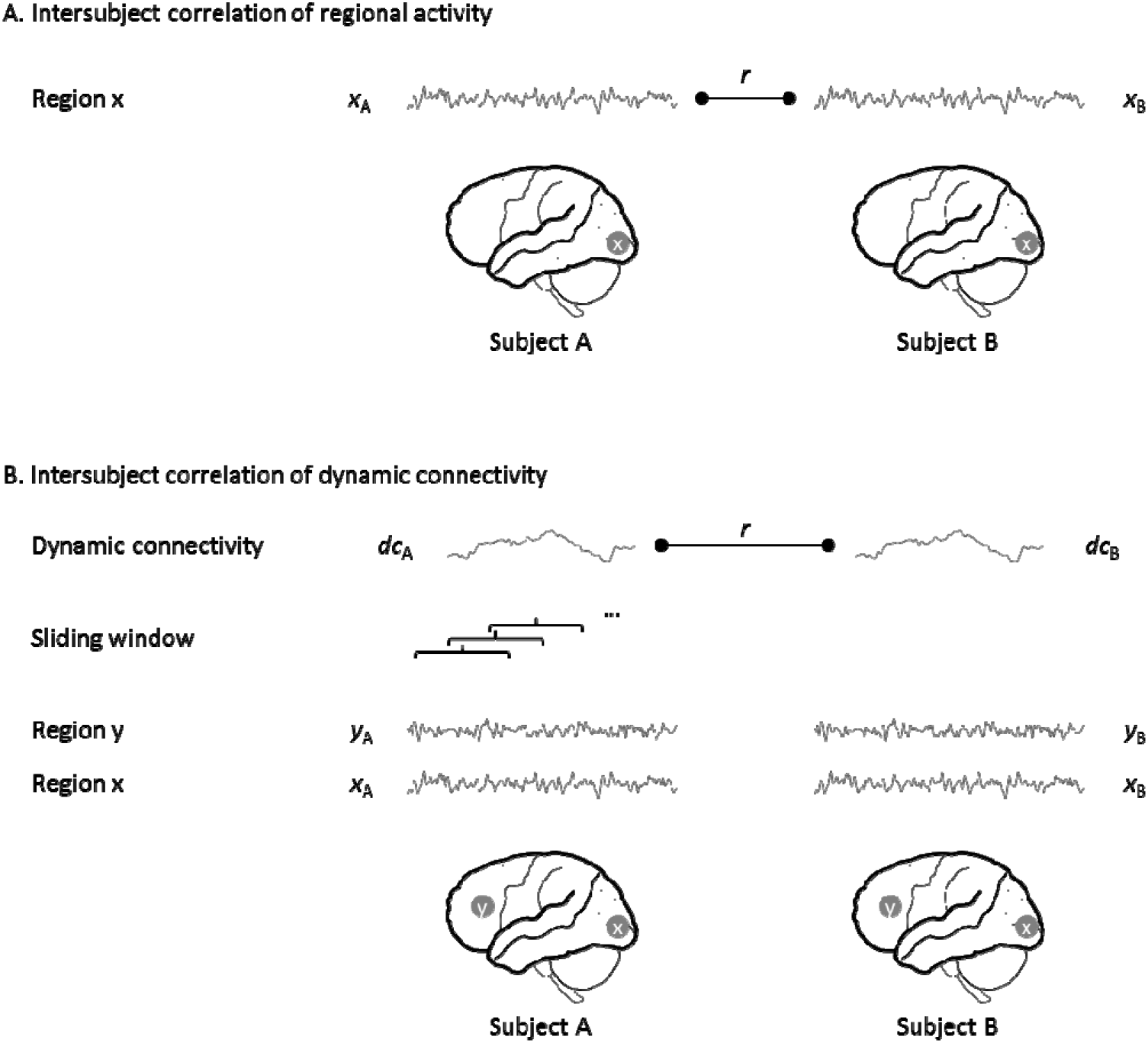
Illustration of the calculations of intersubject correlations of the time series of regional activity (A) and the time courses of dynamic connectivity between two regions (B).

In the current study, we analyzed an fMRI dataset where the subjects were scanned when viewing a short animated movie. The aim was to identify dynamic connectivity that were shared cross subjects during the movie watching. In order to do so, we first performed regular intersubject correlation analysis to identify brain regions that showed consistent regional activity. Given these regions, we adopted a seed-based strategy to calculate dynamic connectivity between a seed region and every voxels in the brain. We then evaluated and identified regions whose connectivity with the seed were consistent cross subjects. Even though higher order association regions did not typically show high intersubject correlations of regional activity (Hasson et al., 2004), their functional communications with lower order regions may be consistent across subject following the narrative of the movie. We therefore hypothesized that intersubject correlations of dynamic connectivity may be able to identify more widespread regions and functional dynamics that are associated with the watching of the movie.

## 2. Materials and methods

### 2.1. Data and task

The fMRI data were obtained through openneuro (https://openneuro.org/; accession #: ds000228). Only the data from adult subjects were analyzed. There were originally 33 adult subjects. Two subjects’ data were discarded because of poor brain coverage (subject #: sub-pixar123 and sub-pixar124), and two were discarded due to large head motions (sub-pixar149 and sub-pixar150). As a result, a total of 29 subjects were included in the current analysis (17 females). The mean age is 24.6 years old (18 to 39 years).

During the fMRI scan, the subjects watched a silent version of Pixar animated movie “Partly Cloudy”, which is 5.6 minutes long (https://www.pixar.com/partly-cloudy#partly-cloudy-1). Brain MRI images were acquired on a 3-Tesla Siemens Tim Trio scanner using the standard Siemens 32-channel head coil. Functional images were collected with a gradient-echo EPI sequence sensitive to BOLD contrast in 32 interleaved near-axial slices (EPI factor: 64; TR: 2□s, TE: 30□ms, flip angle: 90°). The voxel size were 3.13□mm isotropic, with 3 subjects with no slice gap and 26 subjects with 10% gap. 168 functional images were acquired for each subject, with four dummy scans collected before the real scans to allow for steady-state magnetization. T1-weighted structural images were collected in 176 interleaved sagittal slices with 1□mm isotropic voxels (GRAPPA parallel imaging, acceleration factor of 3; FOV: 256□mm). For more information about the dataset please refers to (Richardson et al., 2018).

### 2.2. FMRI data analysis

#### 2.2.1. Preprocessing

FMRI data processing and analyses were performed using SPM12 and MATLAB (R2017b) scripts. A subject’s T1 weighted structural image was first segmented into gray matter, white matter, cerebrospinal fluid, and other tissue types, and was normalized into standard Montreal Neurological Institute (MNI) space. The T1 images were then skull stripped based on the segmentation results. Next, all the functional images of a subject were realigned to the first image of the session and coregistered to the skull stripped T1 image of the same subject. Framewise displacement was calculated for the translation and rotation directions for each subject (Di and Biswal, 2015). Subjects who had maximum framewise displacement greater than 1.5 mm or 1.5° were discarded from further analysis. See supplementary materials section S1 for additional analysis on the head motion effects. The functional images were then normalized to MNI space using the parameters obtained from the segmentation step with resampled voxel size of 3 x 3 x 3 mm^3^. The functional images were then spatially smoothed using a Gaussian kernel of 8 mm. Lastly, a voxel-wise general linear model (GLM) was built for each subject to model head motion effects (Friston’s 24-parameter model) (Friston et al., 1996), low frequency drift (1/128 Hz), and constant offset. The residuals of the GLM were saved as a 4-D image series, which were used for further intersubject correlation analysis. The residual time series were all mean centered because of the constant term included in the GLM.

Removing low frequency drifts in BOLD signals is a critical step for dynamic connectivity analysis (Leonardi and Van De Ville, 2015). Leonardi and Van De Ville have suggested a high-pass filter of 1/W Hz to avoid spurious dynamic connectivity caused by low-frequency fluctuations, where W represents the window length in the sliding window analysis. The high-pass filter of 1/128 Hz is the default in the GLM module in SPM. Given the window length of 60 s (30 TR) in the current analysis, we also applied high-pass filtering of 1/64 Hz in a supplementary analysis. The results are very similar to what using the 1 / 128 Hz high-pass filtering (see supplementary materials section S3).

#### 2.2.2. Intersubject correlation analysis

The correlations of time series of either brain activity or dynamic connectivity are calculated between pairs of subjects. If there are N subjects, then there will be N x (N-1) / 2 correlation coefficients. The statistics of these correlations become tricky, because they are calculated from only N subjects, therefore not independent. An alternative approach is leave-one-out (Nastase et al., 2019), where the time series of one hold-out subject were correlated with the averaged time series of the remaining N – 1 subjects. The averaged time series of N – 1 subjects were thought to reflect the consistent component rather than noisy individual’s time series. Therefore, the resulting correlations should be higher than the pair-wise correlations. Another benefit is that this approach estimates one correlation for each subject, making group level statistics easier. Therefore, we adopt the leave-one-out approach in the current analysis.

We first performed intersubject correlation analysis on regional activity time series. The preprocessed BOLD time series were extracted for each voxel and subject in a gray matter mask. For a given voxel, the time series of one subject was held out, and the averaged time series of the remaining subject were calculated. Then the time series of the hold-out subject were correlated with the averaged time series. This process was performed for every voxel and every subject, resulting in one correlation map for one subject. The correlation maps were transformed into Fisher’s z maps. Group level one sample t test was then performed to identify regions whose intersubject correlations were consistently greater than 0. However, the null hypothesis statistical significance testing may not provide much information of the effect size. There may be only small but consistent correlations for each subject, which could give rise to very high statistical significance in a one sample t test. Indeed, when doing such null hypothesis statistical significance testing for intersubject correlation analysis, usually almost all the brain regions will show somehow significant correlations (Chen et al., 2016). We are more interested and focused on the real effect size, i.e. correlation coefficients, in our analysis. We therefore averaged the Fisher’s z maps, and transformed them back into r maps. The continuous r maps were shown in the results section.

We next performed intersubject correlation analysis on dynamic connectivity using a seed-based approach. Given that a set of brain regions showed high intersubject correlations of regional activity, we defined these regions as seeds. We adopted a relatively high threshold of r > 0.45 for the averaged intersubject correlation map of regional activity to isolate four visual related seeds. Two of them were located in the medial and posterior portion of the occipital lobe, which mainly covered the lingual gyrus and calcarine sulcus. The other two seeds were located bilaterally in the middle occipital gyrus and extended to the middle temporal gyrus. We labeled them as left and right medial visual and lateral visual seeds, respectively. In addition, we adopted a relatively low threshold of r > 0.35 to isolate the left and right supramarginal gyrus seeds. The maps of the six seeds are available at: https://identifiers.org/neurovault.collection:6245.

For each seed, we performed voxel-wise correlation analysis, i.e. calculating intersubject correlations of dynamic connectivity between the seed and every voxel in the gray matter mask. For two given time series from a seed and a voxel, we first applied sliding window technique to calculate dynamic connectivity for each subject. The window length was set as 30 time points (60 s) (Nastase et al., 2019), and the time step was set as 2 time point (4 s). Therefore, the time course of dynamic connectivity had 70 window steps. Next, we calculated correlations between the time courses of dynamic connectivity of a given subject with the averaged dynamic connectivity of the remaining subjects for a given voxel. As a result, there was one correlation map for each seed and each subject.

The r maps of correlations of dynamic connectivity were transformed into Fisher’s z maps for group level statistical analysis. Again, we simply calculated an averaged z map for a seed, and transformed it back into r map. In addition, we performed group-level analysis to identify regions that showed different dynamic connectivity patterns with different levels of seeds. Specifically, we calculated contrast images from the Fisher’s z maps for each subject representing the differences between specific levels of seeds compared with the other seeds. For example, we calculated a contrast image using [1, 1, - 0.5, −0.5, −0.5, −0.5] on the six z maps of a subject to define the differences between the two medial visual seeds and the remaining four seeds. The contrast images were entered into a one sample t test GLM using nonparametric statistics in SnPM13 (Statistical NonParametric Mapping, http://warwick.ac.uk/snpm). Resulting clusters were first formed at p < 0.001, and the cluster extend was thresholded using family-wise error (FWE) corrected p < 0.0167 (0.05 / 3). The cluster level FWE threshold (0.0167) was chosen to further account for the three levels of seeds (medial visual, lateral visual, and supramarginal seeds).

In addition to the voxel-based analysis, we also performed region of interest (ROI)-based analysis for in-depth examinations of the dynamic connectivity effects. In addition to the six seeds, we included three more regions that showed different intersubject correlations of dynamic connectivity with different seeds. Specifically, they were the left precentral gyrus that showed higher intersubject correlations of dynamic connectivity with the medial visual seeds, and the posterior cingulate cortex and medial prefrontal cortex that showed higher intersubject correlations of dynamic connectivity with the supramarginal gyrus seeds. The regions were defined based on the statistical significant clusters from the group-level analysis. The maps of the three regions are available at: https://identifiers.org/neurovault.collection:6245. The calculations of intersubject correlations of dynamic connectivity were the same as the seed-based analysis.

The selections of sliding window length is nontrivial (Fu et al., 2014; Zhang et al., 2013). In addition to the 30-TR window length, we also explored other window length of 10 TRs (20 s), 20 TRs (40 s), 40 TRs (80 s), 50 TRs (100 s), and 60 TRs (120 s). For each window length, we calculated intersubject correlations of dynamic connectivity among the 9 ROIs.

#### 2.2.3. Relations with other measures

We first compared the intersubject consistent dynamic connectivity with stable functional connectivity. For each subject, we calculated correlation coefficients across the 9 ROIs, and transformed them into Fisher’s z. Then the z matrices were averaged across the 29 subjects, and transformed back into r values. In addition, we calculated the consistent component of each ROI, i.e. averaging the time series across the 29 subjects. And then one single correlation matrix among the 9 ROIs was calculated.

Given the consistent component of the 9 ROIs, we also calculated dynamic connectivity between pairs of ROIs using the same sliding window approach. The time courses of dynamic connectivity calculated from the consistent component were compared with the averaged dynamic connectivity that was calculated from each subject.

We further asked whether the observed intersubject consistent dynamic connectivity was driven by the consistent component of regional activity, or by the subject-specific idiosyncratic component. To do so, for each ROI, we regressed out the consistent component from each subject’s time series, and calculated dynamic connectivity from the residual time series for each subject. Intersubject correlations of dynamic connectivity calculated from the residual time series were compared with those from the taw time series.

Lastly, we calculated intersubject correlations of regional activity using the same sliding window approach for the 9 ROIs. That is, for each ROI, intersubject correlation was calculated at each window, resulting in a time course of intersubject consistency of regional activity in each of the ROI.

## 3. Results

### 3.1. Intersubject correlations of regional activity

We first calculated intersubject correlations of regional activity for every voxel in the brain during the video watching (Figure 2A). The highest correlations were around 0.5. The major regions that had high intersubject correlations were the visual cortex extending anterior to the fusiform gyrus and middle temporal lobe. The bilateral supramarginal gyrus also showed high intersubject correlations. The bilateral precentral gyrus also showed intersubject correlations, but the effect sizes were much smaller. Figure 2A shows all the voxels with positive correlation values. It is noteworthy that many regions showed very small intersubject correlations, including largely the prefrontal cortex and anterior temporal lobe.

**Figure 2.**
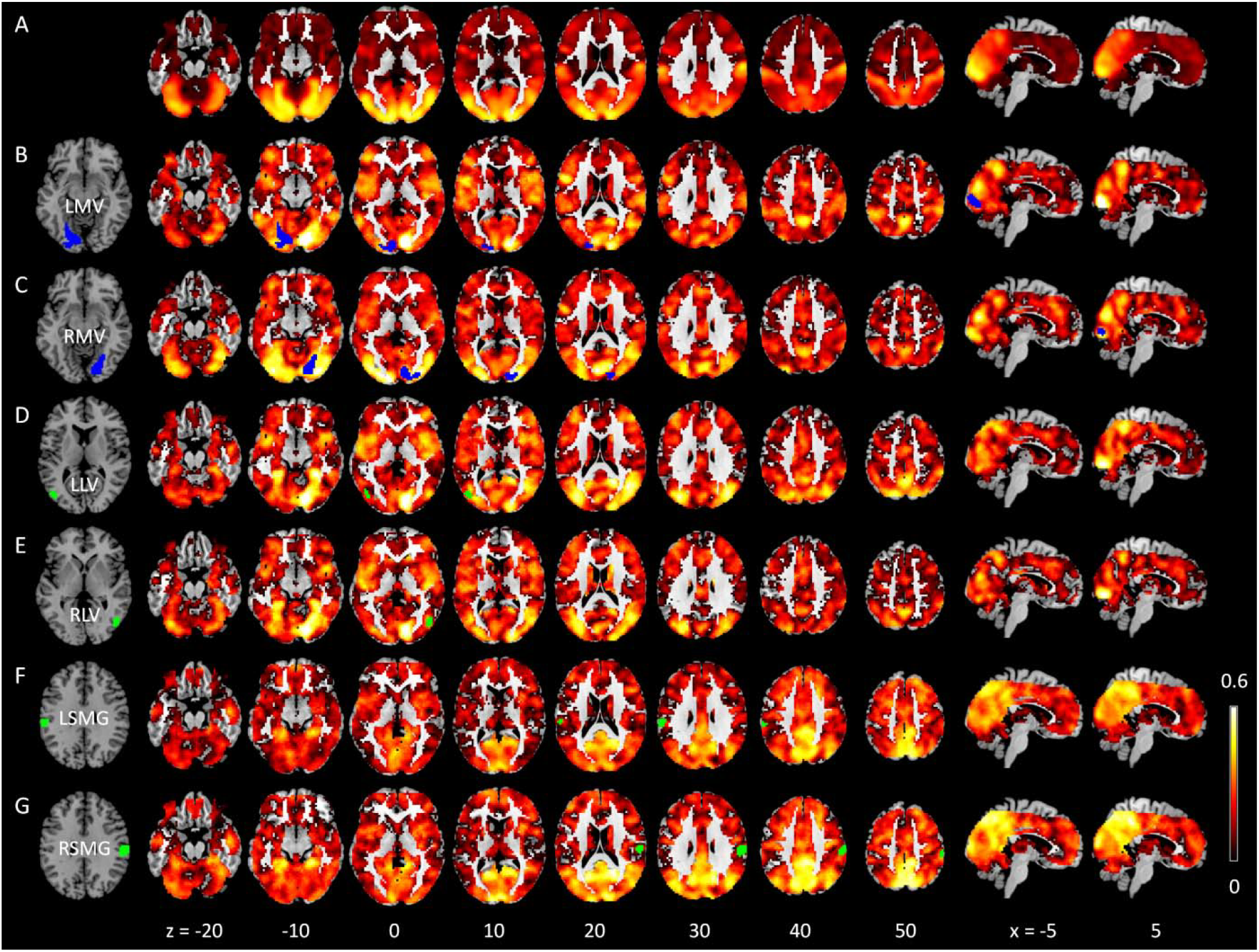
Intersubject correlation maps of regional activity (A) and dynamic connectivity with different seeds (B through G). The seed regions were depicted in blue or green in respective rows. All voxels with positive correlations are shown. The numbers at the bottom represent z or x coordinates in Montreal Neurological Institute (MNI) space. LMV, left medial visual; RMV, right medial visual; LLV, left lateral visual; RLV, right lateral visual; LSMG, left supramarginal gyrus; RSMG, right supramarginal gyrus. All the maps are available at: https://identifiers.org/neurovault.collection:6245.

### 3.2. Intersubject correlations of dynamic connectivity

#### 3.2.1 Seed-based analysis

We defined seed regions where there were high intersubject correlations of regional activity, which included bilateral medial visual regions, lateral visual regions, and supramarginal gyrus. We next calculated voxel-wise intersubject correlations of dynamic connectivity with the six seeds, respectively (Figure 2B through 2G). There were widespread brain regions that showed intersubject consistent dynamic correlations with different seeds. First of all, the effect sizes of the intersubject correlations of dynamic connectivity, i.e. the correlation coefficients, were comparable to those in the intersubject correlations of regional activity. Secondly, regions with intersubject correlations of dynamic connectivity turned out to be more widespread and extended to the frontal and parietal regions that did not show high intersubject correlations of regional activity. See supplementary materials section S2 for direct comparisons between the intersubject correlations of dynamic connectivity and those of regional activity. Thirdly, the left and right corresponding seeds showed similar dynamic connectivity patterns, but there were substantial differences in the patterns of dynamic connectivity among medial visual, lateral visual, and supramarginal gyrus seeds. In order to highlight specific brain regions that showed dynamic connectivity with different seeds, we compared each pair of seeds with the remaining seeds using nonparametric group-level model (Figure 3 and Table 1). The medial visual seeds showed more consistent dynamic connectivity with the left precentral gyrus and occipital regions compared with other seeds. The lateral visual seeds showed more consistent dynamic connectivity with several visual regions compared with the other seeds. In contrast, the supramarginal seeds showed consistent dynamic connectivity with the precuneus/posterior cingulate gyrus and medial prefrontal cortex compared with the other seeds, which basically formed the default mode network.

**Table 1.**
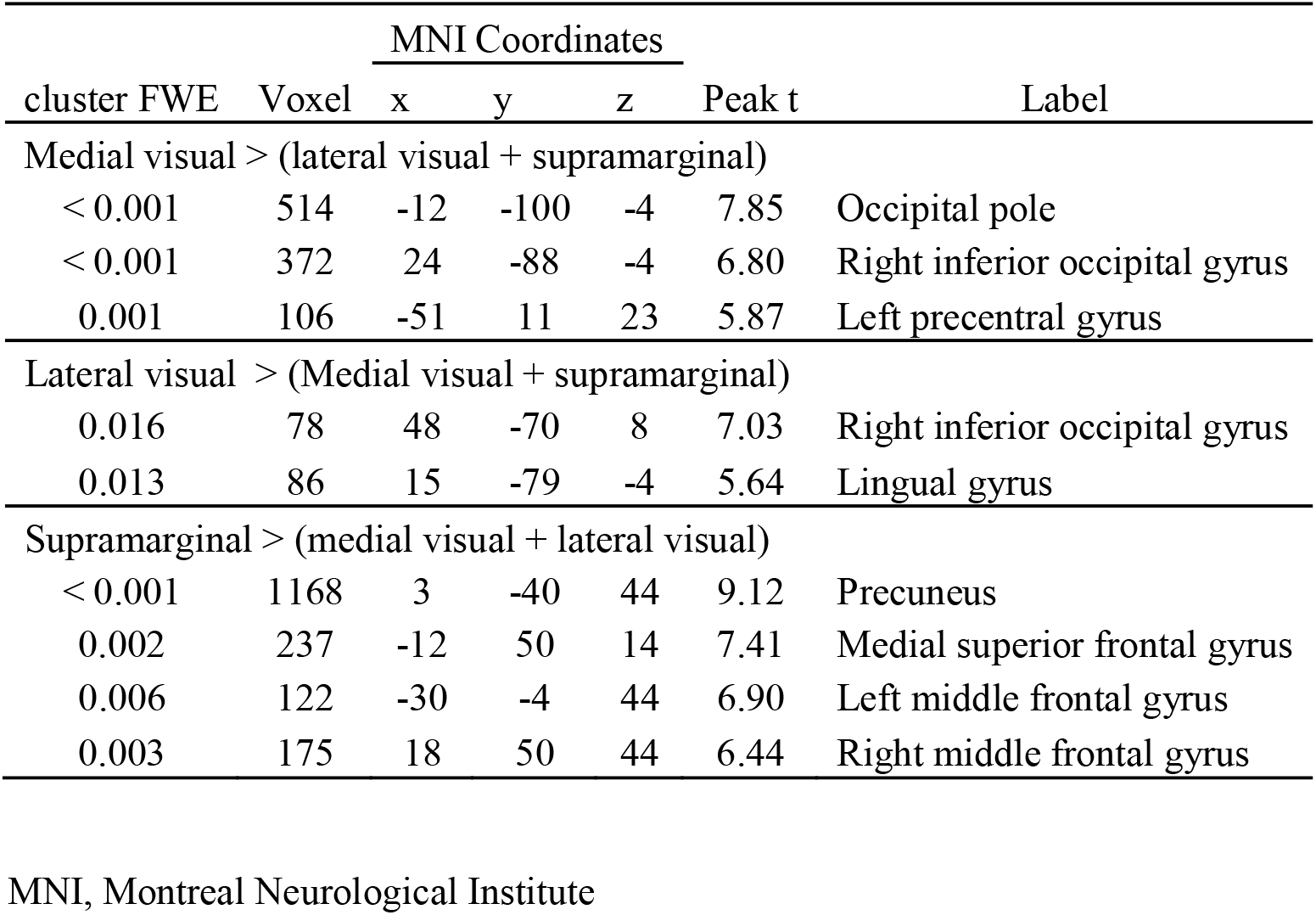
Clusters with differential intersubject correlations of dynamic connectivity among the medial visual, lateral visual, and supramarginal gyrus seeds. All clusters were thresholded at p < 0.001, and cluster thresholded at p < 0.0167 (0.05 / 3) after family-wise error (FWE) correction using nonparametric method.

**Figure 3.**
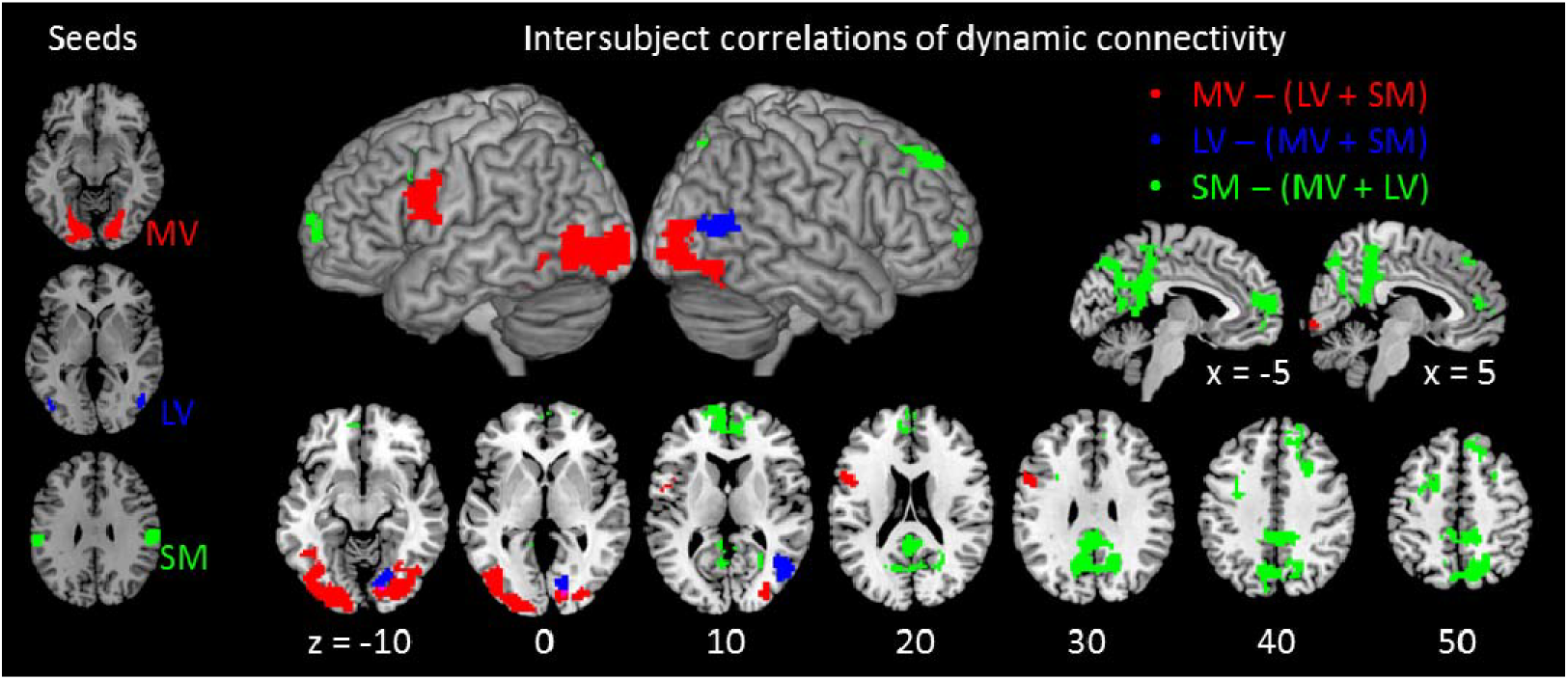
Differential intersubject correlations of dynamic connectivity among the medial visual, lateral visual, and supramarginal gyrus seeds (depicted on the left). All maps were thresholded at p < 0.001, and cluster thresholded at p < 0.0167 (0.05 / 3) after family-wise error (FWE) correction using nonparametric method. MV, medial visual; LV, lateral visual; and SM, supramarginal gyrus. Unthresholded statistical maps are available at: https://identifiers.org/neurovault.collection:6245.

#### 3.2.2. Relations with stable functional connectivity

In order to better understand and interpret the dynamic connectivity and regional functions, we further calculated different types of connectivity measures among a set of regions of interest. In addition to the six seeds, we defined left precentral gyrus, posterior cingulate cortex, and medial prefrontal cortex ROIs that showed different dynamic connectivity with different seeds. Among the 9 regions, we calculated regular mean functional connectivity (Figure 4A) and connectivity derived from the consistent components across the 29 subjects (Figure 4B). These two correlation matrices looked similar, and clearly showed three clusters of brain regions. The first four regions were all visual. The fifth to seventh regions were the bilateral supramarginal gyrus, and lateralized frontal region, which were all high order association brain regions. And the last two regions were part of the default mode network, which showed negative correlations with the association regions in the consistent component correlations.

**Figure 4.**
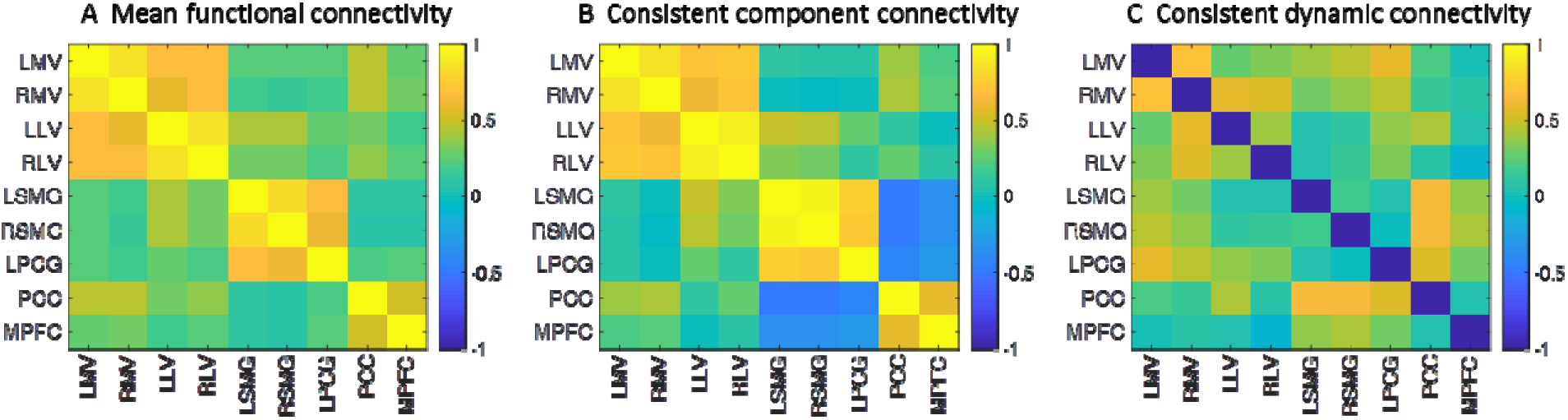
Correlation matrices among the 9 regions of interest (ROI) using different methods. A) Mean functional connectivity across the 29 subjects. B) Correlations of the consistent component of each ROI (averaged time series across the 29 subjects). C) Intersubject correlations of dynamic connectivity. LMV, left medial visual; RMV, right medial visual; LLV, left lateral visual; RLV, right lateral visual; LSMG, left supramarginal gyrus; RSMG, right supramarginal gyrus; LPCG, left precentral gyrus; PCC, posterior cingulate cortex; and MPFC, medial prefrontal cortex.

The intersubject consistent dynamic connectivity matrix (Figure 4C) was largely different from the two stable correlation matrices. Some high consistent dynamic connectivity was observed within the visual regions. The highest correlation was between the left and right medial visual regions (*r = 0.70*). In contrast, many consistent dynamic connectivity were shown between different functional networks, where there were virtually none or even negative stable correlations. Specifically, the medial visual regions showed high consistent dynamic connectivity with the left precentral gyrus ROI. The highest intersubject correlation was 0.56 between left medial visual region and left precentral gyrus. The default mode regions and supramarginal regions also showed high consistent dynamic connectivity. The highest correlation was 0.64 between the posterior cingulate cortex and right supramarginal gyrus. It is noteworthy that these regions generally showed negative stable correlations in Figure 4B.

Lastly, we analyzed the time courses of dynamic connectivity for the above mentioned pairs of regions (Figure 5). The dynamic connectivity between left and right medial visual regions was in general high, which is consistent with the results of stable connectivity. But it can be seen that the connectivity level went down during the first half of windows, and continued with two cycles of up and down fluctuations. The fluctuations rather than a monotonic linear trend suggest that the dynamics of connectivity is not simply due to sensory habituations. The left medial visual region and left precentral gyrus did not show high level of correlations in general. But it had small positive correlations at the beginning of the run, went down to around zero, and then went back to small positive correlations. What is more interesting is the dynamic connectivity between the right supramarginal gyrus and posterior cingulate cortex, where the connectivity switched between positive and negative values during the whole course.

**Figure 5.**
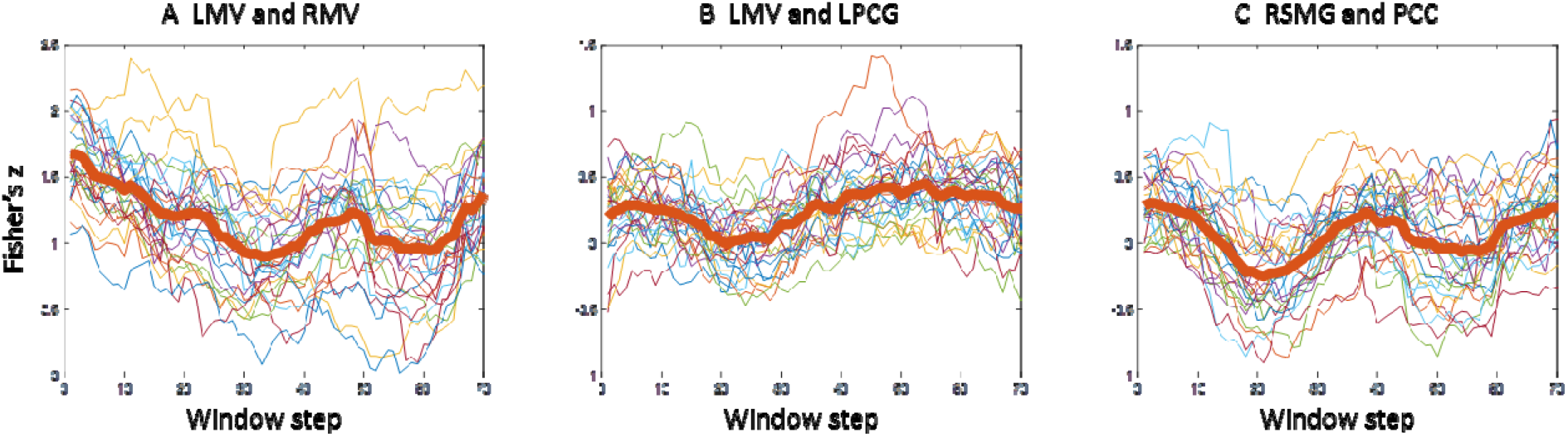
Time courses of dynamic connectivity (Fisher’s z) for three pairs of regions of interest. Each thinner line represents the time course of one subject, and the thicker red lines represent the averaged time courses. LMV, left medial visual; RMV, right medial visual; LPCG, left precentral gyrus; RSMG, right supramarginal gyrus; PCC, posterior cingulate cortex.

#### 3.2.3. Relations with the consistent component

A subsequent question is that whether the observed intersubject consistent dynamic connectivity is driven by the consistent component of regional activity across subject, or by subject-specific idiosyncratic components. We then regressed out the consistent component for each subject’s time series and calculated intersubject correlations of dynamic connectivity from the residual time series (Figure 6B). Compared with the intersubject correlations of dynamic connectivity from the original analysis (Figure 6A), the consistency of dynamic connectivity from the residual time series were largely diminished. Figure 6D and 6E illustrate the changes of dynamic connectivity time courses after the regression between a representative ROI pair, i.e. right supramarginal gyrus and posterior cingulate cortex (see supplementary Figure S4 for other ROI pairs). The intersubject correlation reduced from 0.64 to 0.29. Figure 6C illustrates the dynamic connectivity of the consistent components of regional activity between these two ROIs. The fluctuating pattern was similar to those calculated from individual subject’s original time series (Figure 6D), further confirmed that the consistent dynamic connectivity across individuals was driven by the consistent component of regional activity.

**Figure 6.**
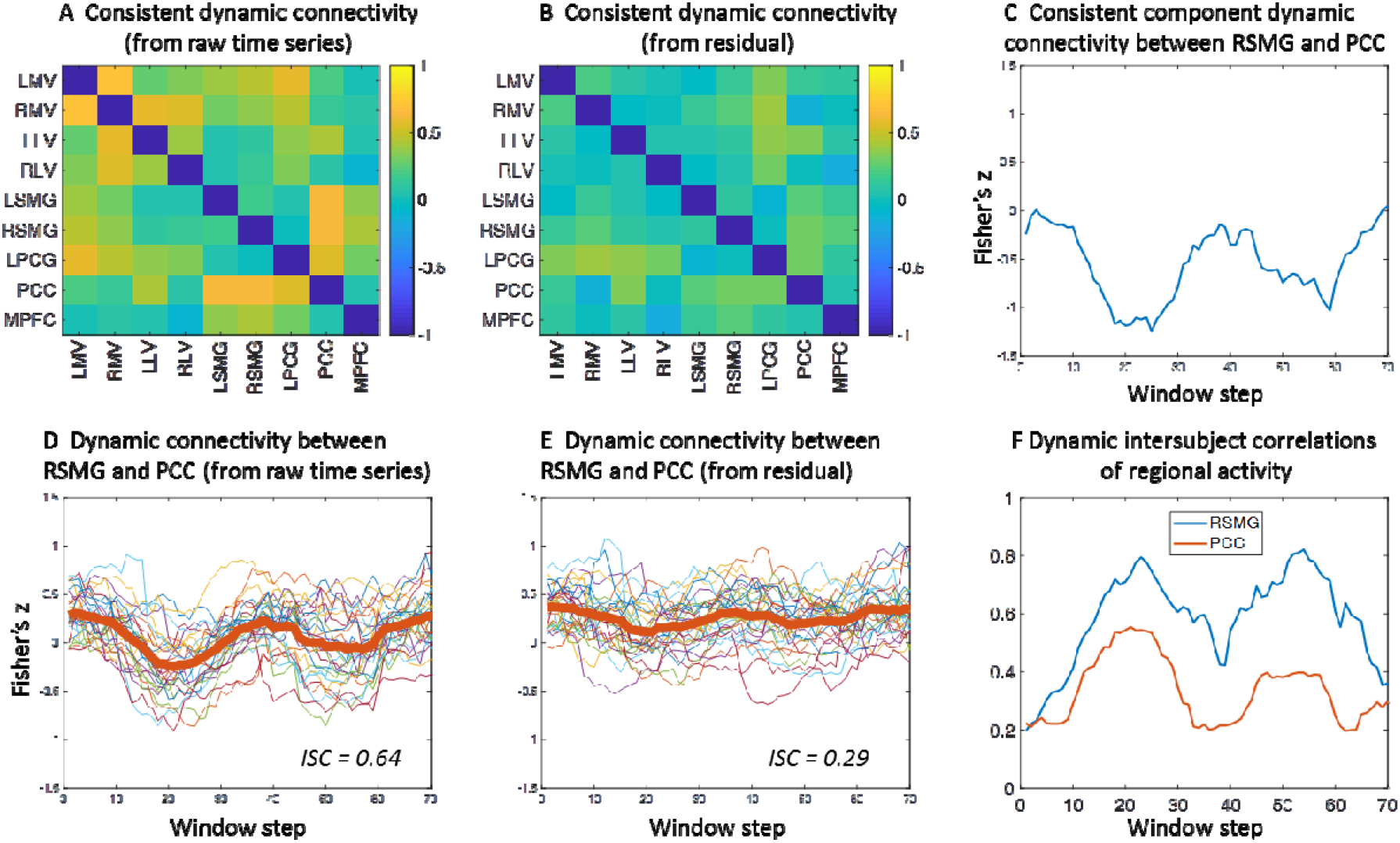
A) and B) Intersubject correlations (ISC) of dynamic connectivity calculated from raw time series (A) and residual time series after regressing out the intersubject consistent components (B). C) Dynamic connectivity of the consistent component of regional activity between right supramarginal gyrus (RSMG) and posterior cingulate cortex (PCC). D) and E) Time courses of dynamic connectivity (Fisher’ z) between RSMG and PCC calculated from raw time series (D) and the residual time series after regressing out the intersubject consistent component. F) The time courses of intersubject correlations of regional activity in RSMG and PCC.

Lastly, we calculated intersubject correlations of regional activity in every sliding window (supplementary Figure S5). Figure 6F shows the time courses of intersubject correlations of regional activity in the right supramaginal gyrus and posterior cingulate cortex ROIs. Both regions showed similarly but reversed time courses as the dynamic connectivity between them. That is, during the two periods of dips of dynamic connectivity, there were elevated intersubject correlations of regional activity in both regions. But this kind of close relations cannot be observed in the other two pairs of ROIs (supplementary Figure S4).

#### 3.2.4.

Effects of sliding-window length We repeated the ROI-based intersubject correlation analysis of dynamic connectivity using different window length from 10 TRs to 60 TRs. The intersubject correlation matrices were in general weaker when the window was shorter, especially for the 10-TR window (Figure 7). As the window went longer, the correlations matrices became similar to the 30-TR window results. But for even longer window, there were two different trends. First, some of the intersubject correlations kept increasing, usually between regions that involved in one or two visual ROIs (Figure 7B). On the other hand, some of the intersubject correlations decreased after peaked at the 30-TR window, usually between regions that involved supramarginal gyrus or posterior cingulate cortex. Figure 7C illustrated the time courses of dynamic connectivity between right supramarginal gyrus and posterior cingulate cortex. It can be seen that the variability of dynamic connectivity time courses were larger in short window. When using 10-TR window, the dynamic connectivity changed fast, and were not aligned across subjects. When using 30-TR window, the dynamic connectivity time courses became smoother, and the fluctuations were more aligned across subjects, which in turn gave rise to higher intersubject correlations. But when using 50-TR window, the time courses of dynamic connectivity become too smooth, so that the subject averaged trend become less apparent. It’s noteworthy that for the dynamic connectivity between the left and right medial visual ROIs and between left medial visual and left precentral gyrus ROIs, there were linear trends of dynamic connectivity across subjects, which gave rise to high intersubject correlations in longer windows (Figure S6).

## 4. Discussion

In the current study, we proposed intersubject correlation analysis on the time courses of dynamic connectivity during natural vision. We were able to identify intersubject consistent dynamic connectivity at similar levels as the intersubject correlations of regional activity, although the time courses of dynamic connectivity were thought to be nosier than the original time series. By using seed regions from the visual cortex and supramarginal gyrus, we demonstrated widespread brain regions that showed high intersubject consistent dynamic connectivity with these seeds, although these regions themselves did not show intersubject correlations of regional activity. These regions included high order association regions such as frontal and parietal regions, as well as the default mode network. The intersubject consistent patterns of dynamic connectivity support the functional meaningfulness of dynamic connectivity during movie watching, and suggest that dynamic connectivity could be a complementary avenue to characterize the functions of a brain region.

**Figure 7.**
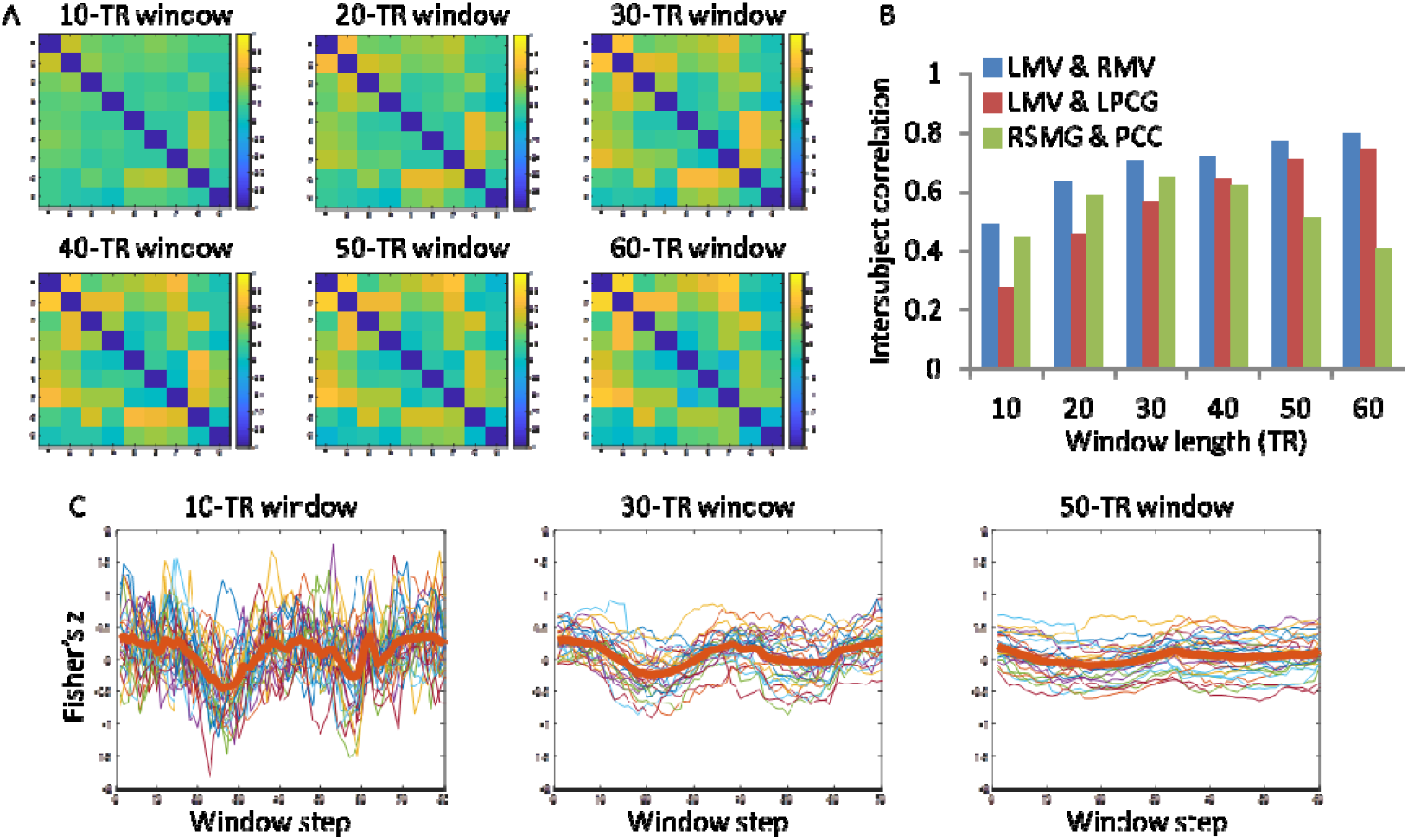
A) The effects of sliding-window length on the intersubject correlations of dynamic connectivity. B) Intersubject correlations of dynamic connectivity of three pairs of regions of interest: left medial visual (LMV) and right medial visual (RMV), LMV and left precentral gyrus (LPCG), and right supramarginal gyrus (RSMG) and posterior cingulate cortex (PCC). C) The time courses of dynamic connectivity between RSMG and PCC in three window lengths. TR, repetition time.

The brain regions that had the highest intersubject correlations of regional activity are mainly in the posterior visual related regions, which are consistent with previous studies (Hasson et al., 2004; Nummenmaa et al., 2012). In addition, the current study found dynamic connectivity among different levels of visual areas that were highly consistent across subjects. This is interesting because although the overall functional connectivity among the visual areas are very high (Figure 4A), there are still functionally meaningful fluctuations of interactions among the visual regions. The observable dynamics of connectivity among visual areas are in line with previous studies showing task modulated connectivity among visual areas in different task conditions (Di et al., 2019, 2015; Di and Biswal, 2017). It is interesting to note that the dynamics of intersubject correlations of regional activity in the visual areas also showed decreased trends at the beginning of the session (Figure S4 and S5). Therefore, the decreased connectivity in the beginning may reflect adaptations effects in the visual areas. However, during the latter half of the session, the dynamics of intersubject correlations of regional activity kept at a stable level, which cannot explain the dynamics of connectivity between them (Figure S4).

The bilateral supramarginal gyrus regions are major regions outside the visual cortex that showed high intersubject correlations of regional activity. The involvements of supramarginal gyrus of intersubject correlations are inconsistent in the literature (Hasson et al., 2004; Kauppi et al., 2010), which probably due to different movies the participants watched. Given their critical role in empathy (Silani et al., 2013), it is reasonable to observe high intersubject correlations in the supramarginal gyrus during the watching of the animated movie, which involves the understanding the intentions of different animated characters. Interestingly, the intersubject correlations of regional activity in the supramarginal gyrus also showed dynamics, with two periods of high correlations roughly between the 20^th^ and 30^th^ windows and between the 50^th^ and 60^th^ windows (Figure 6F and S4B). The first may correspond to the scene when Peck the stork and Gus the cloud first met, where their interactions appeared to be different from the other storks and clouds. The second may coincide with the scene when Peck flew away, and Gus thought Peck had abandoned him. These scenes require active inferences of the intentions of the characters, and may involve mismatches between predictions and the actual story development. Therefore, it is reasonable to see high cross-subject similarities in the supramarginal gyrus during these two periods.

In addition to regional activity, we also found that the default mode network showed highly consistent dynamic connectivity with the supramarginal gyrus regions. Similar to the supramarginal gyrus ROIs, the regional activity in the posterior cingulate cortex showed two periods of high consistent regional activity (Figure 6F). But interestingly, during these two periods the posterior cingulate cortex and supramarginal gyrus showed strong anti-correlation (Figure 6C). The default mode network involves high-order representation of the world (Carhart-Harris and Friston, 2010). And the functional communications between the default mode network and supramarginal gyrus may reflect the prediction error between inner representation and the input from the video. Similar to a previous study using dynamic intersubject connectivity analysis (Simony et al., 2016), both of the studies highlighted the critical role of the default mode network in understanding of the narratives of a movie.

Generally speaking, the intersubject consistent connectivity and stable functional connectivity showed disassociations. Specifically, the ROI pairs that showed highly consistent dynamic connectivity may have high stable functional connectivity or very low overall connectivity. The latter case may be more interesting, because it suggests transient functional communications that cannot be observed in traditional functional connectivity analysis. The 9 ROIs used in the current analysis are from three functional modules, i.g. unimodal visual, higher order task positive, and default mode networks. The three functional modules can be confirmed in the matrix of stable connectivity (Figure 4A), where there are high within-module functional connectivity but weak between-module connectivity. The matrix of consistent dynamic connectivity, on the other hand, showed that there are more between-module dynamic connectivity. These results are in line with the economy account of brain network organizations, which suggests that the functional communications between modulates are costly in terms of energy consumption, therefore are more transient (Bullmore and Sporns, 2012). It is also consistent with the findings that the connectivity between modules are more variable and context dependent (Di and Biswal, 2019; Fu et al., 2017).

By calculating dynamic connectivity time courses from individual’s time series, the proposed method focused on the consistency of the dynamic connectivity time courses across subjects. Our additional analysis showed that the consistent dynamic connectivity time courses was driven by the dynamic connectivity of the consistent component of the regional time series, at least for the current video watched. The latter method provides a simple approach to reveal the dynamics of connectivity, and is closely related to the dynamic intersubject functional connectivity approach proposed by Simony et al. (Simony et al., 2016). Our method, on the other hand, can not only reveal the time course of dynamic connectivity, but can also provide a quantity of a connection about how the dynamic connectivity is consistent across subjects. Eventually, we will be able to obtain a matrix of the consistency of dynamic connectivity among ROIs from the whole brain. This is important because the seed-based approach used in the current analysis may miss dynamic connectivity between regions that do not have consistent regional activity. The connectome-based approach can provide a comprehensive mapping of dynamic communications across the brain during the watching of a movie, and can be seen a special form of task connectome (Di and Biswal, 2019).

The selection of window length for dynamic connectivity analysis is nontrivial (Fu et al., 2014; Zhang et al., 2013). The shorter the window length, the finer the temporal resolution for dynamic connectivity could be. However, less time points for each window would also mean noisier estimates of connectivity. In the current analysis, 10-TR window gave very noisy estimate of functional connectivity, thus making intersubject correlations very low. On the other hand, longer window will make connectivity estimate accurate, but at a cost of losing temporal resolution. In the context of the current video watched, 30-TR window seems a balance. However, this time scale of dynamic connectivity fluctuations may not be easily generalized to other videos or to resting-state. But it certainly can provide some insight to the chosen of window length in future studies. In addition, some computational method may be used to avoid the window length issue, e.g. using adaptive covariance estimates (Fu et al., 2014; Zhang et al., 2013) or window-free method such as Kalman filtering (Kang et al., 2011) or instantaneous phase synchronization (Glerean et al., 2012).

## 5. Conclusion

In the current study, we proposed intersubject correlation analysis on dynamic connectivity. The results revealed widespread brain regions that showed consistent intersubject correlations of dynamic connectivity. The consistent correlations support the functional significance of dynamic connectivity during natural vision. The method may provide a complementary approach to understand the dynamic nature of brain functional integrations.

## Acknowledgement

This study was supported by grants from (US) National Institute of Health (R01 AT009829; R01 DA038895).

