## Supplementary materials for "Intersubject consistent dynamic connectivity during natural vision revealed by functional MRI"

**Contents:**

S1. Head motion effects

S2. Higher intersubject correlations of dynamic connectivity than those of regional activity

S3. Effects of high-pass filtering

S4. Relations with the consistent component

S5. Effects of sliding-window length

**S1. Head motion effects**

We calculated framewise displacement for translation (Figure S1A) and rotation (Figure S1D) separately for all the subjects (Di and Biswal, 2015). It can be seen that there is not much synchronized head motion across all the subjects. This can be confirmed by calculating correlations matrices of the framewise displacement time series across the subjects (Figure S1B and S1E). The distributions of all the intersubject corelations are plotted in Figure S1C and S1F. The mean and median of the correlations were around 0 for both directions. In addition, we also calculated intersubject correlations of framewise displacement using the leave-one-out method. The averaged correlations were 0.045 and 0.042 for translation and rotation, respectively. The intersubject correlations of head motion were much smaller in amplitude than the intersubject correlations of regional activity and dynamic connectivity reported in the main text. Therefore, head motion is not likely to contribute to the observed intersubject correlations in regional activity and dynamic connectivity.

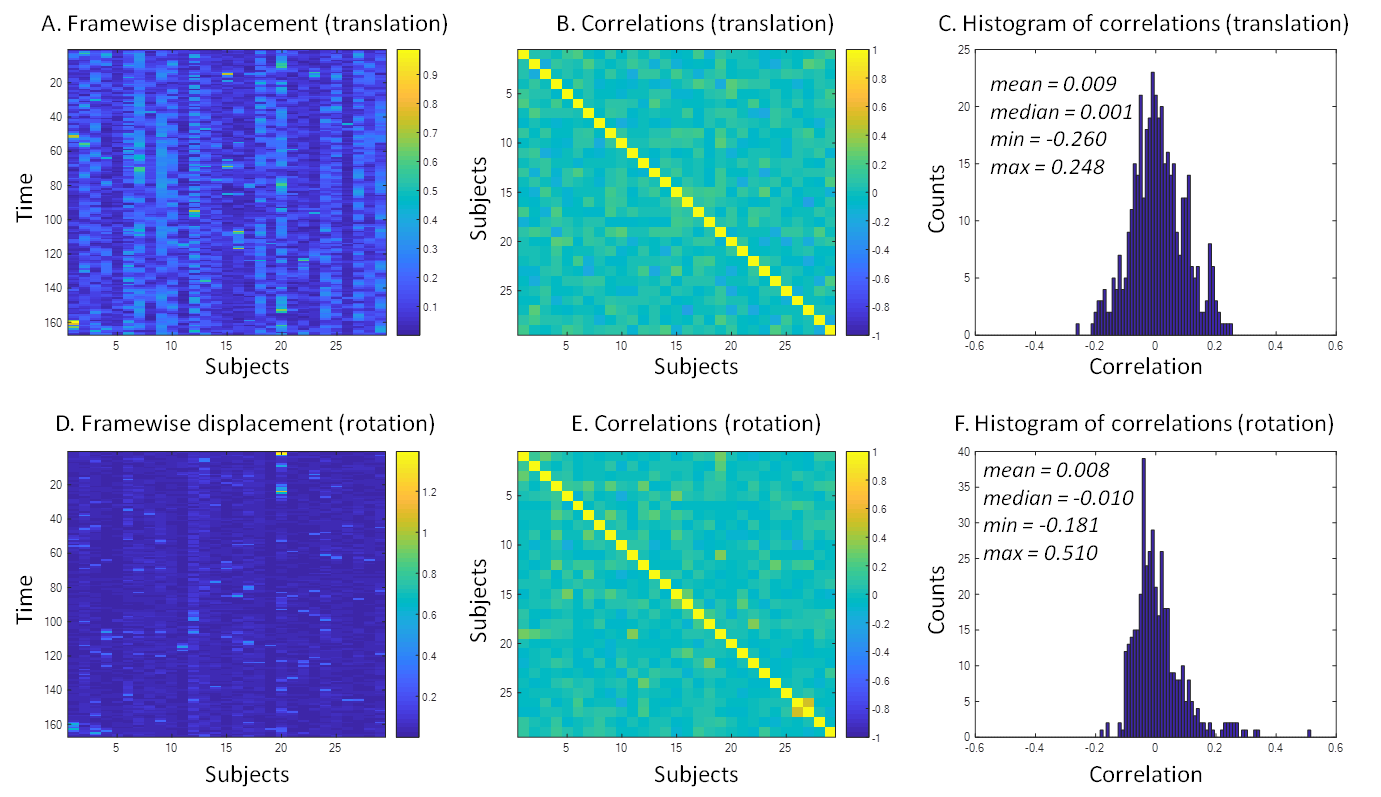

**Figure S1** Intersubject correlations of head motion. The head motion effects were quantified as framewise displacement in translation (upper row) and rotation (lower row). A and D demonstrate the time courses of framewise displacement of all the 29 subjects. The correlations of the time courses of framewise displacement across the 29 subjects are shown in B and E. The histograms of the pairwise correlations of head motion are shown in C and F.

**S2. Higher intersubject correlations of dynamic connectivity than those of regional activity**

We directly compared the intersubject correlations calculated from dynamic connectivity with those from regional activity. Group-level analysis was performed using paired t test general linear model (GLM), with nonparametric statistics using SnPM13 (Statistical NonParametric Mapping, <http://warwick.ac.uk/snpm>). We are only interested in higher correlations for dynamic connectivity than regional activity. A one-tailed cluster forming threshold of p < 0.001 was used, and the cluster-level inference was done using family-wise error (FWE) correction of p < 0.05/6 to additionally account for the six seeds. Statistical significant clusters are shown in Figure S2. Four out of the six analyses revealed significant effects. Consistent with our observations from the raw maps, widespread regions, including frontal lobe, insula, and parietal regions showed greater intersubject correlations with different seeds than the intersubject correlations of their own regional activity.

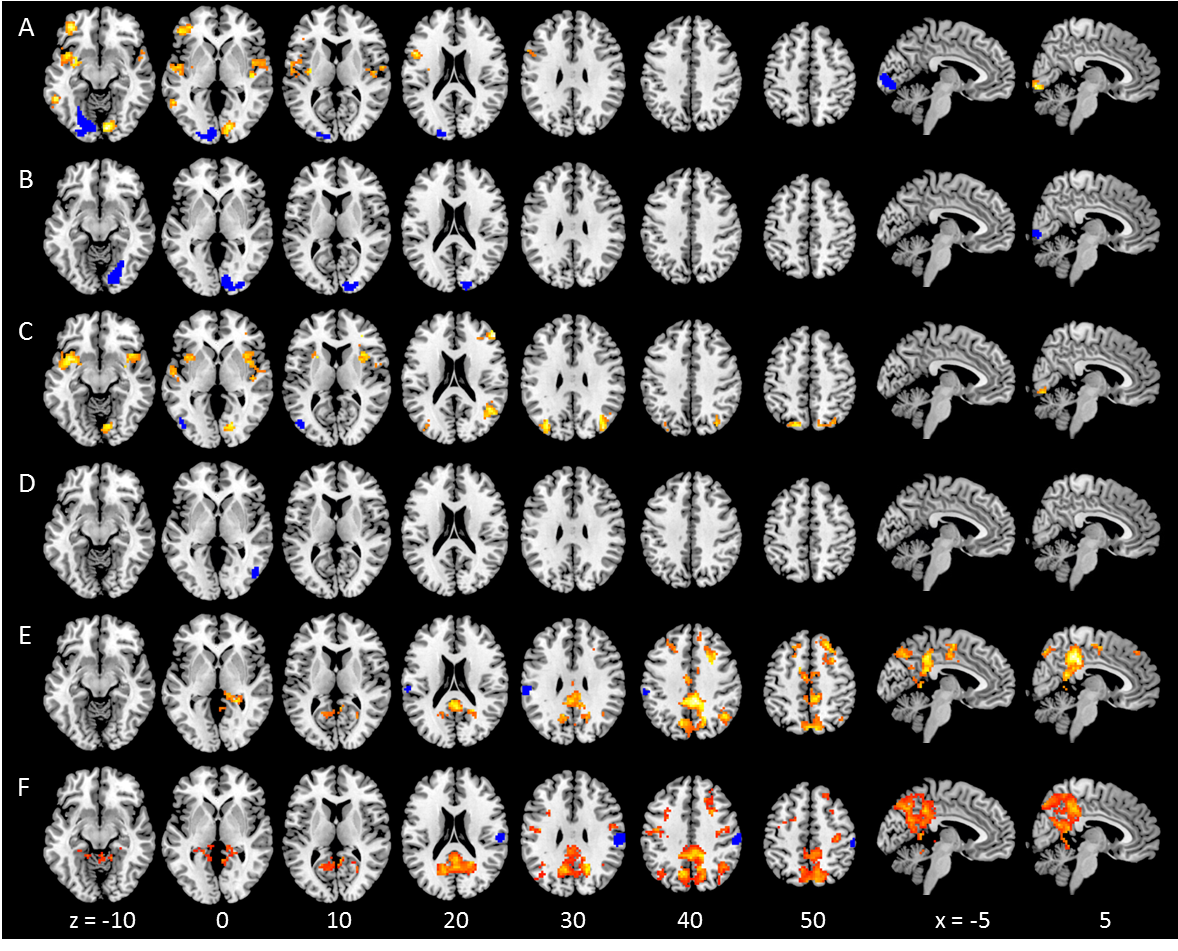

**Figure S2** Clusters with higher intersubject correlations of dynamic connectivity than those of regional activity. The seed regions were depicted in blue in respective rows. The clusters were first formed using a one-tailed threshold of p < 0.001, and the cluster-level inference was done using nonparametric method using family-wise error (FWE) correction of p < 0.05/6 to additionally account for the six seeds. The numbers at the bottom represent z or x coordinates in Montreal Neurological Institute (MNI) space.

**S3. Effects of high-pass filtering**

In this analysis, we applied high-pass filter at 1/128 Hz through general linear modeling in SPM. But a study has suggested a high-pass filter of 1/W Hz to avoid spurious dynamic connectivity caused by low-frequency fluctuations, where W represents the window length of the sliding window method (Leonardi and Van De Ville, 2015). Given the window length of 60 s (30 TR) in the main analysis, we re-run the analysis by using a high-pass filter at 1/64 Hz, and repeated the ROI-to-ROI dynamic connectivity analysis. The resulting matrix of consistent dynamic connectivity (bottom row of Figure S3) looked very similar to the original matrix. We also plotted individual dynamic connectivity in the three representative pairs of ROIs. In all the three cases the intersubject correlation values reduced when using the 1/64 Hz filter compared with when using the 1/128 Hz filter, but the amount of changes were very limited.

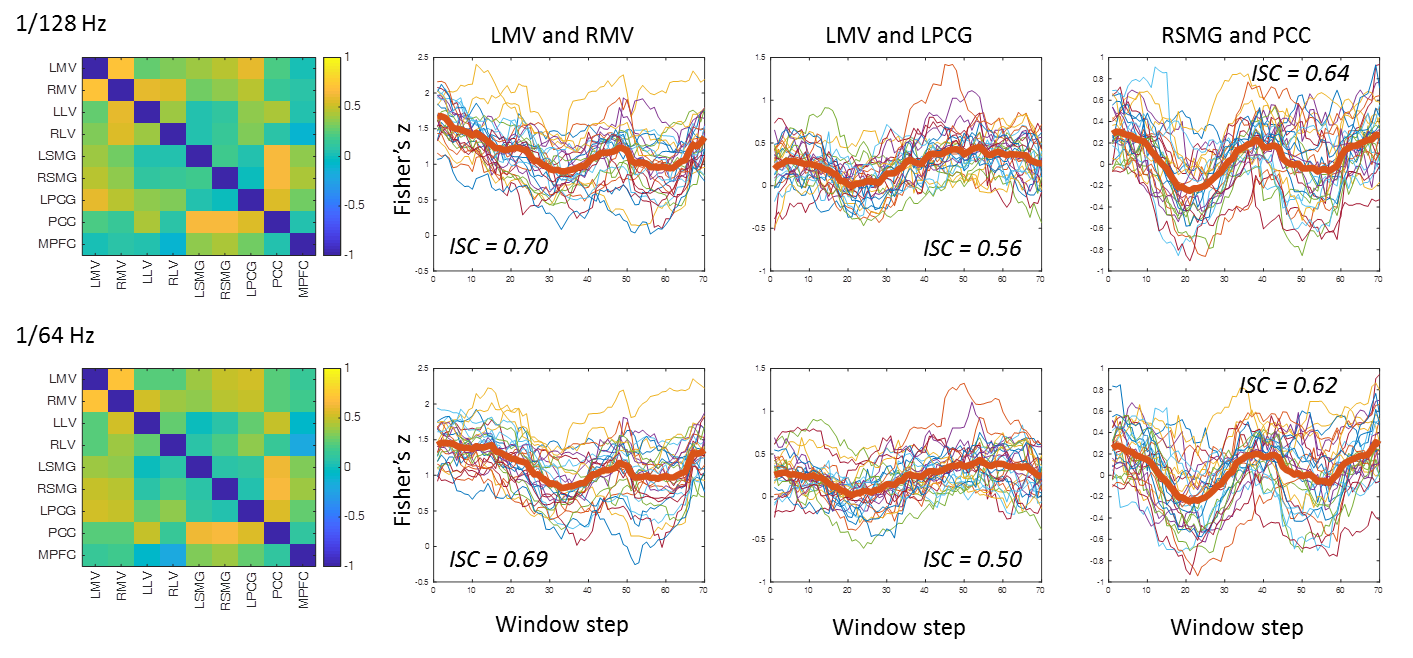

**Figure S3** Top row, intersubject correlation of dynamic connectivity results in the original analysis using a high-pass filtering of 1/128 Hz in the preprocessing. Bottom row, intersubject correlation of dynamic connectivity results using a high-pass filtering of 1/64 Hz in the preprocessing. LMV, left medial visual; RMV, right medial visual; LPCG, left precentral gyrus; RSMG, right supramarginal gyrus; PCC, posterior cingulate cortex; ISC, intersubject correlation.

**S4. Relations with the consistent component**

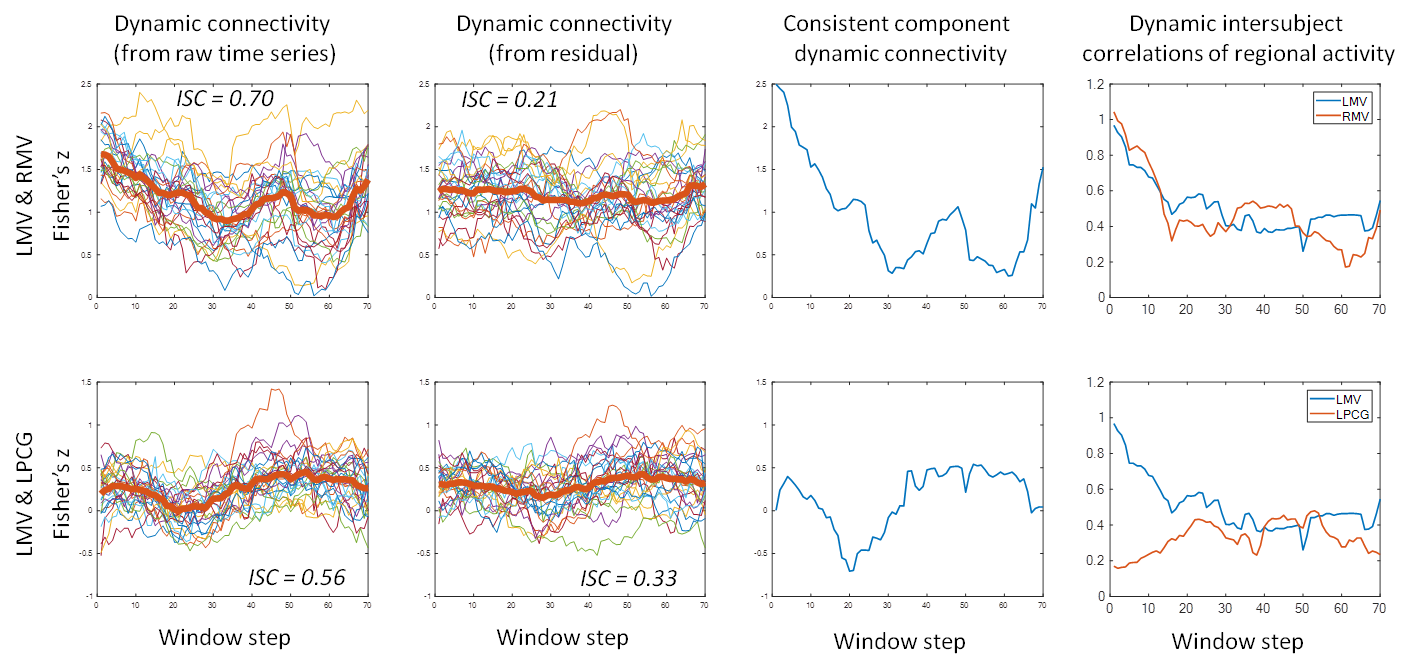

**Figure S4** Top row, functional connectivity and regional activity in the left and right medial visual ROIs (LMV and RMV). Bottom row, functional connectivity and regional activity in the left medial visual and left precentral gyrus (LPCG) ROIs. The left most column, time courses of dynamic connectivity (Fisher’s z) calculated from raw time series. The second left column, time courses of dynamic connectivity (Fisher’s z) calculated from the residual time series after regressing out the intersubject consistent component. The second right column, dynamic connectivity of the intersubject consistent components of regional activity. The right most column, the time courses of intersubject correlations of regional activity. ISC, intersubject correlation.

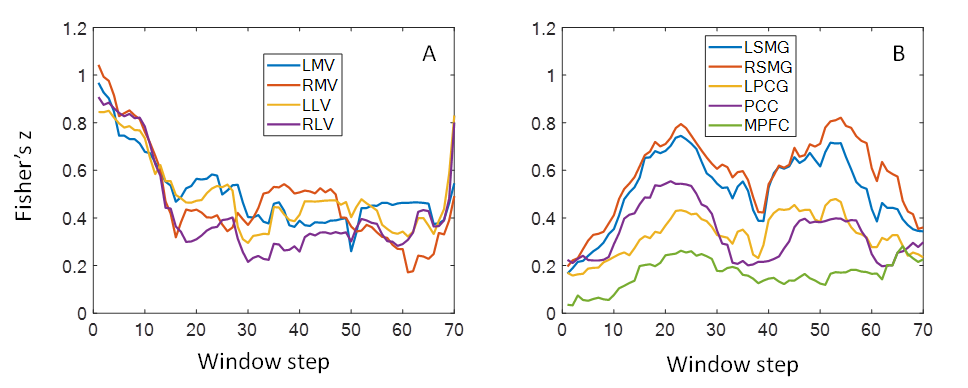

**Figure S5** Dynamic intersubject correlations of regional activity of all the nine ROIs. These regions showed two distinct patterns. A, the four visual ROIs had very high intersubject correlations at the beginning of the session, and decreased to a relative stable level around 0.4 (A). On the other hand, the remaining five ROIs showed similar trends (B), which all looked like a reversed version of the dynamic connectivity between right supramarginal gyrus and posterior cingulate cortex ROIs. Among them, the left and right supramarginal gyrus ROIs had the strongest intersubject correlations at the two peaks around 20 and 50 window steps.

**S5. Effects of sliding-window length**

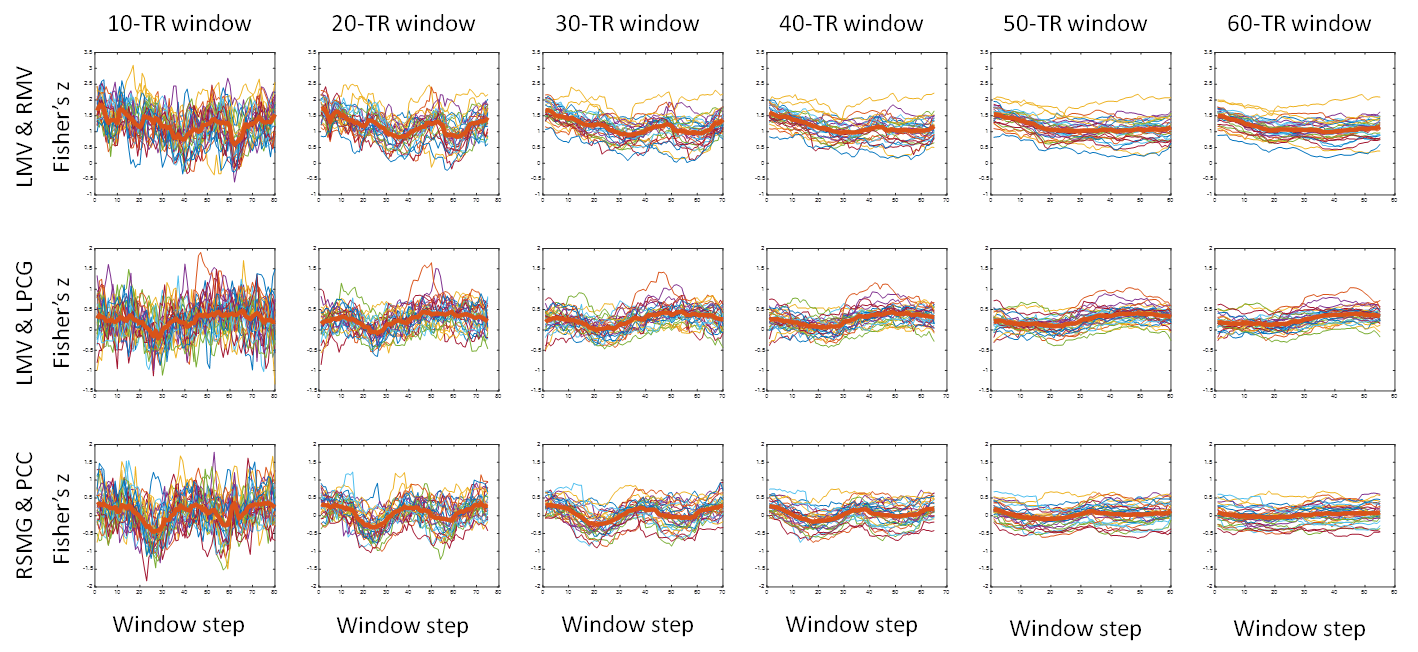

**Figure S6** The effects of sliding-window length on the time courses of dynamic connectivity between three pairs of regions of interest. Upper row, left medial visual (LMV) and right medial visual (RMV); middle row, left medial visual and left precentral gyrus (LPCG); and bottom row, right supramarginal gyrus (RSMG) and posterior cingulate cortex (PCC).
